# Incubation of discriminative stimulus-controlled cocaine craving: An animal model relevant to relapse prevention

**DOI:** 10.1101/494070

**Authors:** Rajtarun Madangopal, Brendan J. Tunstall, Lauren E. Komer, Sophia J. Weber, Jennifer K. Hoots, Veronica A. Lennon, Jennifer M. Bossert, David H. Epstein, Yavin Shaham, Bruce T. Hope

## Abstract

In abstinent drug addicts, cues formerly associated with drug-taking experiences gain relapse-inducing potency (“incubate”) over time. Animal models of incubation may help develop treatments to prevent relapse, but these models have ubiquitously focused on the role of conditioned stimuli (CSs) signaling drug delivery. From a translational perspective this is problematic because people encounter these stimuli only *during or after* relapse. For this reason, incubation in response to discriminative stimuli (DSs) that signal drug availability *before* relapse, not yet examined in preclinical studies, could be more relevant to relapse prevention. We trained rats to self-administer cocaine (or palatable food) under DS control, then investigated DS-controlled incubation of craving, in the absence of drug-paired CSs. DS-controlled cocaine (but not palatable food) seeking incubated over 60 days of abstinence and persisted up to 300 days. Understanding the neural mechanisms of this DS-controlled incubation holds significant promise for drug relapse treatments.

## Introduction

The risk of relapse is a major obstacle for effective treatment of drug addiction (*1*, *2*). In abstinent drug users, several factors contribute to drug relapse, including exposure to cues and contexts previously associated with drug use (*3*), stressors (*4*), or acute exposure to the drug itself (*5*). Preclinical studies have recapitulated these effects in relapse models using mice, rats, and nonhuman primates (*6*, *7*). A major finding across these studies is that cue-induced drug-seeking (in the absence of the drug) increases progressively during abstinence, a phenomenon termed *incubation of drug craving* (*8*, *9*). Time-dependent increases in drug-seeking have been demonstrated in rats trained to self-administer cocaine, heroin, methamphetamine, alcohol, and nicotine, as well as non-drug rewards such as sucrose (*10*). These findings in rodents mirror incubation of cue-induced drug craving and physiological responses in human addicts (*11*–*14*), and have been important for studying neural mechanisms contributing to drug relapse (*15*).

Preclinical incubation models have shown how cues presented after performance of a drug-taking response and paired with subsequent drug delivery during training potentiate drug-seeking when presented response-contingently during abstinence. These “confirmatory” conditioned stimuli (CSs) inform the laboratory animal that the drug-taking response has been completed during training. From a translational perspective, this is problematic: when an abstinent addict encounters this kind of stimulus (e.g., the sensation of vapor inhaled into the lungs or the early interoceptive effects of the drug), relapse will have already happened. Early preclinical studies of incubation showed that it could also occur in the absence of discrete drug-paired CSs (*8*, *16*). This suggests that incubation could also be induced by other stimuli associated with drug-taking, such as the contextual cues (e.g. the chamber used for operant training) or discriminative stimuli (DSs) that signal drug availability (e.g., the house-light that is illuminated during the training session, the retractable lever that serves as the operant manipulandum). Surprisingly little is known about this CS-independent incubation. A recent study suggested that it is not mediated by contextual cues (*17*), leaving DSs as a likely culprit. DSs are different from cues typically investigated in these studies in that they are neither response-contingent like CSs, nor ever-present like contextual cues. Rather, DSs signal drug availability—or unavailability—thereby preceding and guiding the performance of drug-taking behavior. Previous studies have shown that a DS signaling drug availability (DS+) can promote persistent drug-seeking behavior while a DS signaling drug unavailability (DS-) can inhibit drug-taking behavior and drug-priming-induced reinstatement of drug seeking (*18*–*26*).

In this study, we sought to directly assess the contribution of DSs to incubation, in the absence of drug-paired CSs. To this end, we first designed a trial-based procedure to train male and female rats to discriminatively self-administer cocaine (0.75 mg/kg/infusion) during trials in which a DS+ signaled cocaine availability, and to suppress responding on the same lever during trials in which a DS-signaled cocaine unavailability during the same session. Drug infusions were not paired with CSs. We then tested for the ability of DSs to control cocaine seeking at multiple time points extending up to 400 days of abstinence. Further, after complete cessation of cocaine-seeking behavior, we assessed whether a priming dose of cocaine would reinstate DS-controlled cocaine seeking in the same rats. Finally, to determine whether DS-controlled incubation under our experimental conditions was specific to cocaine, we trained a separate group of rats on an analogous procedure using palatable food (45 mg high-carbohydrate pellets) as the operant reward and assessed the time course of DS-controlled food-seeking behavior.

## Results

### Experiment 1: Incubation of discriminative-stimulus-controlled cocaine seeking

#### Training

The experimental timeline and individual trial design are shown in Fig. 1A,B. Rats learned to respond on the lever for cocaine reward during the first six sessions of continuous access (Fig. 1C). They continued responding in the trial format and then learned to discriminate DS+ from DS-during discrimination training. The number of “successful” trials (denoted as *trials* and defined as making at least one lever press during a trial) and total number of lever presses (denoted as *lever presses* and recorded separately for each DS trial type) during each session were analyzed. We used a two-way maximum-likelihood-based multilevel model with within-subjects factors Session (discrimination sessions 5-14) and DS (DS+, DS-). For *trials*, we observed a significant main effect of DS (F_1,13_=948.21, p<0.0001) but not of Session, and no interaction between Session and DS, indicating that responding during DS+ trials was higher than responding during DS- trials during all the discrimination training sessions. For *lever presses*, we observed a significant main effect of DS (F_1,13_= 161.63, p<0.0001), and an interaction between Session and DS (F_9,117_=2.62, p=0.0085) but no main effect of Session. Post-hoc analyses indicated that responding during DS+ trials was higher than responding in DS-trials during the last four discrimination training sessions (p<0.05).

**Figure 1.**
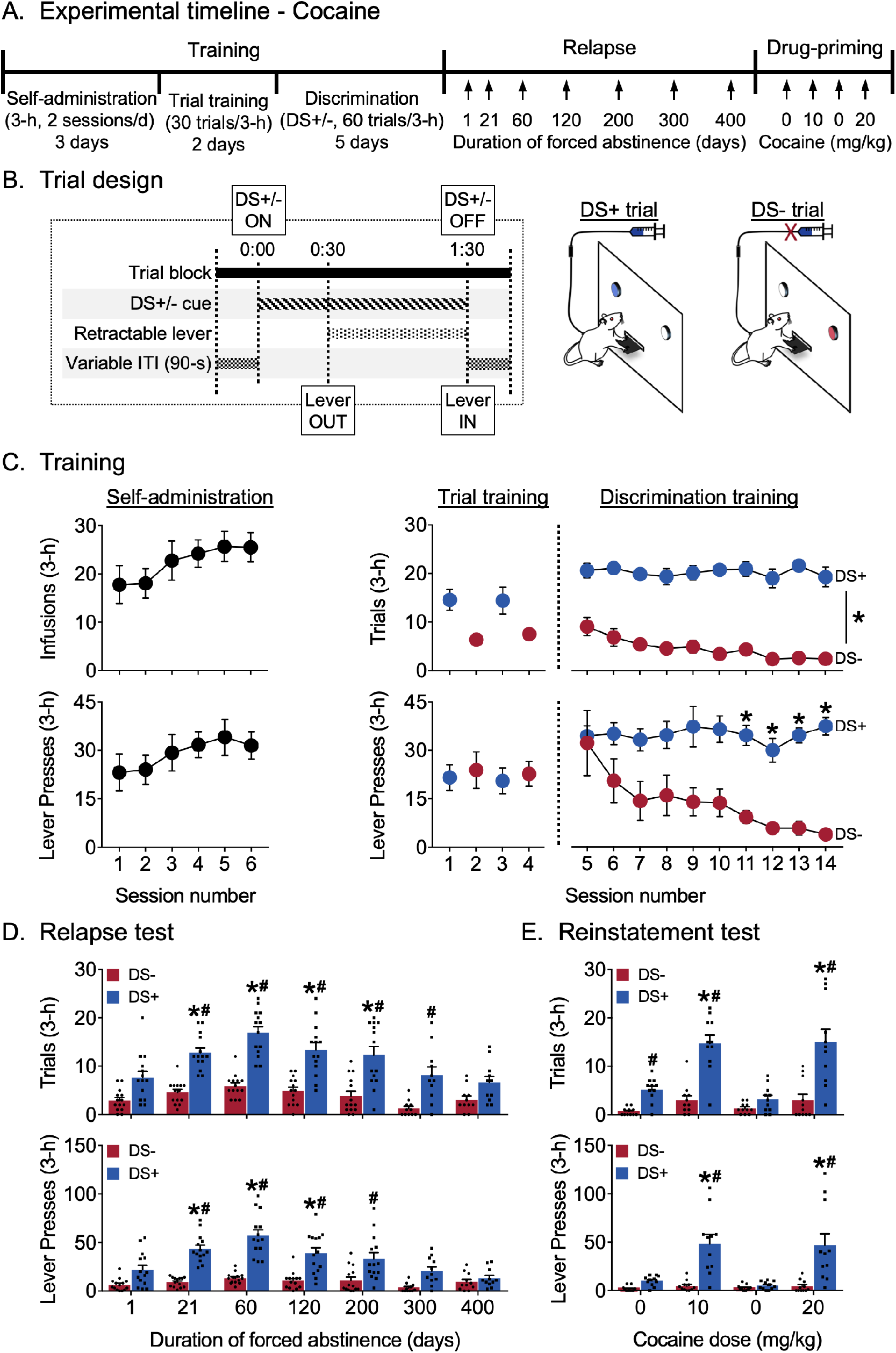
Incubation of discriminative-stimulus-controlled cocaine seeking. (A) Experimental timeline. (B) Schematic showing the timing of individual events during a single 3-minute DS trial, and the differences between the two trial types during discrimination training for cocaine reward. Rats received cocaine reward (0.75 mg/kg/infusion) during DS+ trials but did not receive cocaine reward during DS-trials (n=16). (C) Training data. *<u>Selfadministration:</u>* Rats learned to self-administer cocaine over 6 sessions. Mean (±SEM) number of cocaine infusions and lever presses during each 3-h session. *<u>Trial training:</u>* Mean (±SEM) number of DS+ or DS-trials with at least one lever press (denoted as *trials*), and number of lever presses during the 3-h sessions (denoted as *lever presses*) with 30 trials of a single trial type (DS+ trials in the AM session, DS-trials in the PM session). *<u>Discrimination training:</u>* Over 10 sessions, rats learned to discriminate DS+ from DS-trials. Mean (±SEM) number of *trials* and *lever presses* during the 3-h discrimination training session with 30 trials each of DS+ and DS-trials presented in a pseudorandomized manner. *indicates significant difference (p<0.05) between responding during DS+ and DS-trial types (n=14). (D) Relapse test. Incubation of lever responding during DS+, but not DS-, trials peaked at 60 days of abstinence and returned to basal levels over 400 days. Mean (±SEM) number of *trials* and *lever presses* during the 3-h relapse test sessions (30 trials each of DS+ and DS-presented in a pseudorandomized manner) under extinction conditions. *denotes significant (p<0.05) difference from responding during day 1. Columns indicate mean (±SEM) for the group, while dots indicate values for individual rats. #denotes significant (p<0.05) difference between DS+ and DS-responding during the test sessions (n=11-14). (E) Reinstatement test. Rats reinstated DS-controlled cocaine-seeking in response to IP injections of cocaine (10 and 20 mg/kg), but not saline. Mean (±SEM) number of *trials* and *lever presses* during the 3-h saline- or cocaine-primed reinstatement test sessions (30 trials each of DS+ and DS-presented in a pseudorandomized manner) under extinction conditions. Columns indicate mean (±SEM) for the group, while dots indicate values for individual rats. *denotes significant (p<0.05) difference from responding on the first saline-prime test session (cocaine dose = 0 mg/kg). #denotes significant difference (p<0.05) between DS+ and DS-responding during the test session (n=11).

#### Relapse test

Figure 1D shows relapse in terms of mean responding during 3-hour non-reinforced discrimination test sessions for cocaine seeking. The same rats were tested at different time points 1-400 days following discrimination training. As during training, we analyzed both *trials* and *lever presses* measures. We used a two-way factorial model with within-subjects factors of Days of forced abstinence (1, 21, 60, 120, 200, 300, and 400 days), and DS type (DS+, DS-). For *trials*, we observed significant main effects of Day (F_6,72_=9.68, p<0.0001) and DS (F_1,13_=257.53, p<0.0001), and an interaction between the two (F_6,72_=4.30, p=0.0009), reflecting higher numbers of “successful” trials associated with DS+ presentation after 21, 60, 120 and 200 abstinence days compared to that at 1 abstinence day (p<0.05), and more “successful” DS+ trials compared to DS-trials on abstinence days 21, 60, 120, 200 and 300 (p<0.05). The number of “successful” DS-trials did not significantly increase over days. For *lever presses*, we observed significant main effects of Day (F_6,72_=8.94, p<0.0001) and DS (F_1,13_=182.25, p<0.0001), and an interaction between the two factors (F_6,72_=7.95, p<0.0001), reflecting higher numbers of lever presses associated with DS+ presentation after 21, 60, and 120 abstinence days compared to that at 1 abstinence day (p<0.05), and higher responding during DS+ trials compared to DS-trials on abstinence days 21, 60, 120 and 200 (p<0.05). The number of lever presses during DS-trials did not significantly increase over days. Overall, the *trial* data indicate incubation of “successful” trials during DS+, but not DS-, trials after 21-200 days of abstinence, while the *lever presses* data indicate incubation of *the number* of lever presses during DS+, but not DS-, trials after 21-120 days abstinence. Further, the rats maintained discriminative responding up to 300 days (by the *trials* measure) or 200 days (by the *lever presses* measure) after the last training session.

#### Reinstatement test

Figure 1E shows reinstatement in terms of mean responding during a 3-hour non-reinforced discrimination session for cocaine seeking after priming injections of either cocaine or saline. For both *trials* and *lever presses* measures, we used a two-way factorial model with the within-subjects factors Treatment condition (Saline 1, 10 mg/kg cocaine, Saline 2, 20 mg/kg cocaine) and DS type (DS+, DS-). For *trials*, there were significant main effects of Treatment (F_3,30_=15.35, p<0.0001) and DS (F_1,10_=108.66, p<0.0001), and an interaction between the two factors (F_3,30_=12.42, p<0.0001), reflecting higher numbers of “successful” trials associated with DS+ presentation after both cocaine priming doses (10 and 20 mg/kg) compared to saline, and higher responding during DS+ trials compared to DS-trials following both cocaine priming doses as well as after the first saline priming injection (p<0.05). The number of “successful” trials associated with DS-presentation was not altered by the Treatment conditions. For *lever presses*, we observed significant main effects of Treatment (F_3,30_=8.31, p=0.0004) and DS (F_1,10_=45.73, p<0.0001), and an interaction between the two factors (F_3,30_=9.45, p=0.0001), reflecting higher numbers of lever presses during DS+ trials after both cocaine-priming injections (10 and 20 mg/kg) compared to saline (Saline1 and Saline2), and higher responding during DS+ trials compared to DS-trials during both cocaine-priming injections (p<0.05). The number of lever presses associated with DS-presentation was not altered by the Treatment conditions. Overall, the *trials* and *lever presses* data indicated reliable cocaine-primed reinstatement during DS+, but not DS-, trials that occurred more than 400 days after the last discrimination training session.

### Experiment 2: Abatement of discriminative-stimulus-controlled palatable food-seeking

#### Training

Rats learned to respond on the lever for palatable food reward during the first three continuous access sessions (Fig. 2C). The rats continued responding in the trial format and then learned to discriminate DS+ from DS-during discrimination training. For analysis of successful discrimination on both *trials* and *lever presses* measures, we used a two-way factorial model with within-subjects factors Session (discrimination training sessions 3-13) and DS (DS+, DS-). For *trials*, we observed significant main effects of Session (F_10,140_=15.42, p<0.0001) and DS (F_1,14_=1014.94, p<0.0001), and an interaction between the two factors (F_10,140_=8.22, p<0.0001), reflecting higher responding during DS+ trials for all but the first session (p<0.05). For *lever presses*, we observed significant main effects of Session (F_10,140_=15.63, p<0.0001) and DS (F_1,14_=577.71, p<0.0001), and an interaction between the two factors (F_10,140_=2.31, p=0.0151), again reflecting higher responding during DS+ trials for all but the first session (p<0.05).

**Figure 2.**
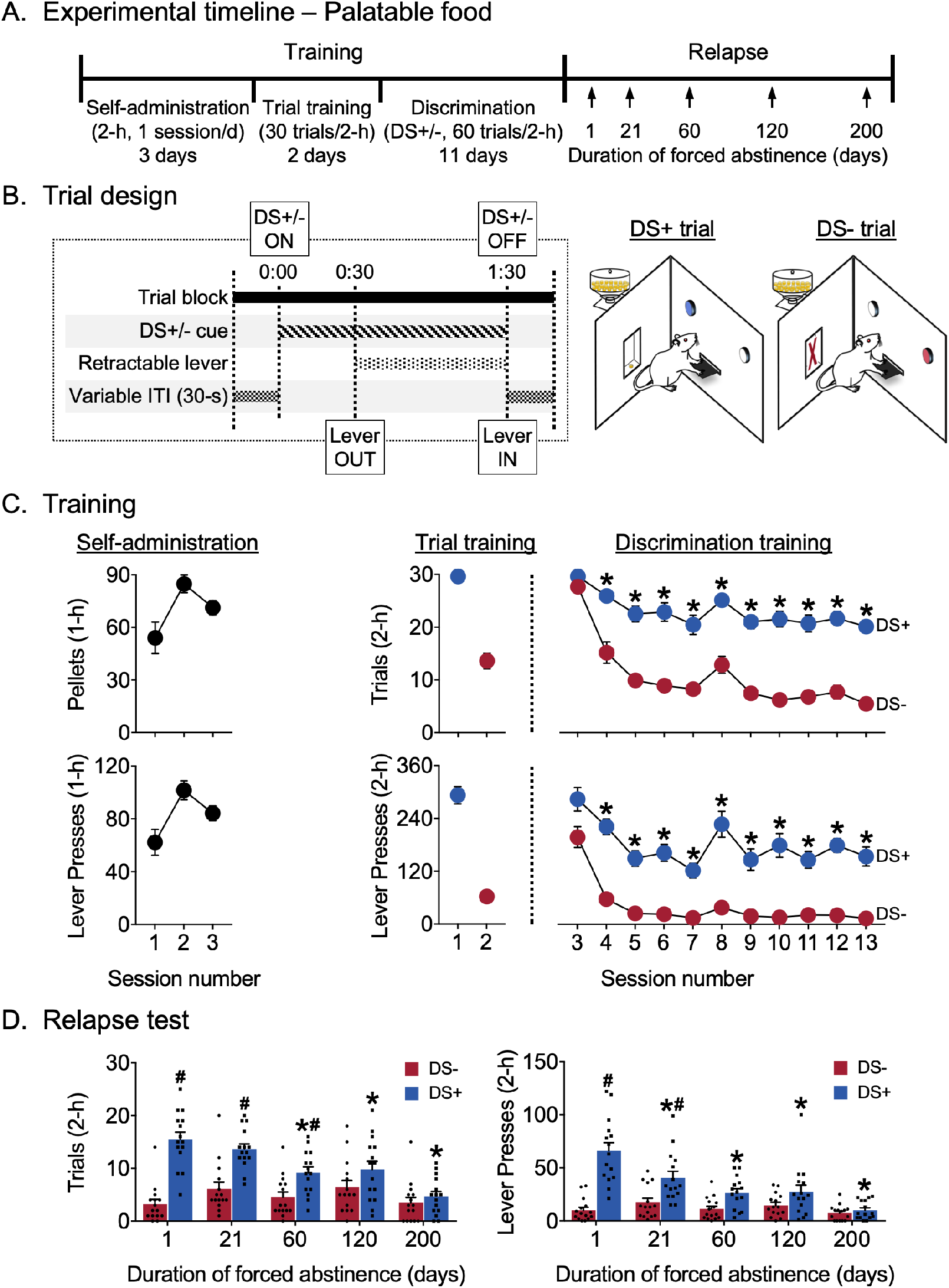
Reduction of discriminative-stimulus-controlled palatable food seeking. (A) Experimental timeline. (B) Schematic showing the timing of individual events during a single 2-minute DS trial, and the differences between the two trial types during discrimination training for palatable food reward (45 mg high carbohydrate pellets). Rats received food reward during DS+ trials but did not receive reward during DS-trials (n=16). (C) Training data. *<u>Selfadministration:</u>* Rats learned to self-administer palatable food over 3 sessions. Mean (±SEM) number of palatable food pellets received and lever presses during each 1-h session. *<u>Trial training:</u>* Mean (±SEM) number of DS+ or DS-trials with at least one lever press (denoted as *trials*), and number of lever presses during the 2-h sessions (denoted as *lever presses*) with 30 trials of a single trial type (DS+ trials in the AM session, DS-trials in the PM session). *Discrimination training:* Over 11 sessions, rats learned to discriminate DS+ from DS-trials. Mean (±SEM) number of *trials* and *lever presses* during the 2-h discrimination training session with 30 trials each of DS+ and DS-trials presented in a pseudorandomized manner. *indicates significant difference (p<0.05) between responding during DS+ and DS-trials (n=15). (D) Relapse test. Lever responding during DS+, but not DS-, trials peaked at 1 day of abstinence and abated over 200 days. Mean (±SEM) number of *trials* and *lever presses* during the 2-h relapse test sessions (30 trials each of DS+ and DS-presented in a pseudorandomized manner) under extinction conditions. *denotes significant (p<0.05) difference from responding during day 1. Columns indicate mean (±SEM) for the group, while dots indicate values for individual rats. #denotes significant (p<0.05) difference between DS+ and DS-responding during the test (n=15).

#### Relapse test

Figure 2D shows relapse in terms of mean responding during 2-hour non-reinforced discrimination test sessions for palatable food seeking. The same rats were tested at different time points 1-200 days following discrimination training. As during training, we analyzed both *trials* and *lever presses* measures. For both *trials* and *lever presses* measures, we used a two-way factorial model with within-subjects factors of Days of forced abstinence (1, 21, 60, 120, and 200 days) and DS type (DS+, DS-). For *trials*, we observed significant main effects of Day (F_4,56_=5.57, p=0.0008), DS (F_1,14_=133.04, p<0.0001), and an interaction between the two factors (F_4,56_=14.29, p<0.0001), reflecting lower numbers of “successful” trials associated with DS+ presentation after 60, 120 and 200 abstinence days than after 1 abstinence day (p<0.05), and higher responding to DS+ than DS-on abstinence days 1, 21, and 60 (p<0.05). The number of “successful” trials associated with DS-presentation did not increase over days. For lever presses, we observed significant main effects of abstinence Day (F_4,56_=8.57, p<0.0001) and DS (F_1,14_=101.92, p<0.0001), and an interaction between the two (F_4,56_=17.44, p<0.0001), reflecting lower numbers of lever presses associated with DS+ presentation after 21, 60, 120 and 200 abstinence days than after 1 abstinence day (p<0.05), and higher responding to DS+ than DS-on abstinence days 1 and 21 (p<0.05). The number of lever presses associated with DS-presentation did not change over days. Overall, *trials* and *lever presses* data indicate that food seeking decreased or abated over time, and that the rats maintained discriminative responding for only 60 days (by the *trials* measure) or 21 days (by the *lever presses* measure) after the last training session.

## Discussion

We used a trial-based procedure to investigate incubation of cocaine or palatable-food seeking controlled by discriminative stimuli that signal availability (DS+) or unavailability (DS-) of the rewards, in the absence of reward-paired CSs. Rats readily learned to respond to the DS+ for either cocaine (experiment 1) or food (experiment 2) and to inhibit responding to the DS-within the same session. DS-controlled cocaine seeking was maximal after 60 days of abstinence (reflecting incubation of DS-controlled cocaine-seeking) and persisted for up to 300 days. Additionally, when DS-controlled cocaine seeking was fully extinguished after 400 days of abstinence, priming injections of cocaine reinstated cocaine seeking. In contrast, DS-controlled food seeking was maximal at 1 day of abstinence, progressively decreased over time, and was no longer observed after 60 abstinence days. Thus, incubation of DS-controlled reward seeking under our experimental conditions was specific for cocaine.

In previous studies, DSs paired with cocaine self-administration have been shown to promote drug seeking that is highly resistant to extinction across multiple non-reinforced test sessions (*21*, *23*, *25*). Reward deliveries in these studies were paired with additional discrete CSs and the contrasting DSs were paired with different levers and presented in separate sessions, making it difficult to disentangle the potential contribution of DSs from CSs and contextual stimuli. In our experiments, reward deliveries were not paired with additional CSs during training, and the two contrasting DSs were paired with a common retractable lever and presented in a pseudo-randomized order within the same session. Following training, the rats were tested under non-reinforced conditions for DS-controlled drug seeking, using the same DS presentation schedule as during training. Because the same operant manipulandum and response was required to seek reinforcement in response to each DS, within the same test session, we know that discriminated drug seeking in our model was exclusively controlled by the DSs and not by contextual stimuli, classically conditioned spatial cues, presentation of the operant manipulandum, or even performance of the drug-seeking response. Under these conditions, we observed persistent nonreinforced drug seeking during DS+ presentations but not DS-presentations, up to 300 days after the last DS-drug pairing. These data extend previous studies of DS-controlled drug-seeking (in the presence of drug-paired CSs) and suggest that in addition to setting the occasion for drug-seeking behavior, the DS+ can acquire excitatory motivational properties.

We demonstrate that DS-controlled cocaine-seeking potentiates during abstinence (that is, we show incubation of DS-controlled cocaine craving) even in the absence of explicit drug-paired CSs. Incubation studies have typically employed between-subjects testing procedures in which rats previously trained to self-administer an addictive drug are returned to the same chambers after varying periods of abstinence and tested for drug-seeking during presentation of the drug-paired CSs (*10*, *27*, *28*). Under these testing conditions, incubation has also been observed in the absence of drug-paired CSs (*8*, *16*). However, the factors controlling this CS-independent incubation phenomenon were never elucidated. In the present study, we found a time-dependent increase in drug seeking (incubation) during DS+ presentations, in the absence of any drug-paired CSs, suggesting that the cocaine-DS+ in our procedure acquired excitatory motivational properties and this property of the DS+ could underlie CS-independent incubation during abstinence. The incubation of DS-controlled cocaine seeking is even more remarkable when considering the within-subjects design used in the present experiments. DS-controlled seeking continued to increase up to 60 days into abstinence and persist up to 300 days (over half the lifespan of a rat) despite exposure of the same group animals to repeated relapse tests under extinction conditions. It is possible that DS-controlled incubation may persist even longer in the absence of extinction learning over repeated relapse tests.

In contrast, cocaine seeking in DS-trials did not incubate – rats continued to suppress responding in DS-trials during all relapse tests and maintained discrimination out to 300 days of abstinence. DSs signaling cocaine unavailability have been shown to inhibit ongoing cocaine self-administration and to suppress cocaine-priming-induced reinstatement (*24*). From the perspective of translation and treatment development, the inhibition of cocaine seeking may be just as important as its potentiation. The behavior guided by each DS in our study was able to survive multiple extinction sessions, and during subsequent tests of cocaine-priming-induced reinstatement – after DS+ responding was extinguished to DS-levels – priming injections of cocaine reinstated cocaine seeking specifically during DS+, but not DS-, trials. Future studies with this procedure will dissociate the neurobiological mechanisms that allow these two functionally orthogonal DSs to mediate incubation of DS-controlled cocaine seeking.

Using a similar format of DS and lever presentation, we also trained rats to lever press for palatable food reward during DS+, but not DS-, trials. We found that food-DS rats made more total responses than cocaine-DS rats during training, maintained their discrimination responding under nonreinforced conditions, and also showed higher seeking responses than cocaine-DS rats during the initial relapse test on day 1. However, under the same repeated-testing schedule used for cocaine relapse, they quickly extinguished their DS-controlled responding in the absence of food and progressively decreased food seeking during abstinence. It is possible that DS-controlled food seeking would have incubated in the absence of repeated relapse testing in extinction. Incubation has been observed using the classical procedure with oral sucrose reward, but the effect was somewhat weaker and not as persistent as incubation of cocaine craving (*8*). The superior strength and persistence of seeking in response to drug-over food-DSs observed here agrees with earlier studies directly comparing drug and food paired-stimuli (*23*, *25*, *29*, *30*). Future studies are required to determine whether this divergence of DS effects on drug versus food seeking is due to differences in the strength of the initial DS-reward associations during training or due to drug-specific neuroadaptations that emerge during abstinence (*15*, *31*).

Taken together, the results of the present experiments show that DS-controlled operant drug seeking but not food seeking incubates during prolonged abstinence. As we noted above, DS-controlled behaviors offer an especially promising path to treatment development because DSs are always present before and during human drug taking; they do not merely accompany or follow it. They can play a critical role in relapse; for example, a study measuring flight attendants’ cigarette craving showed that craving peaked toward the end of flights as the opportunity to smoke a cigarette neared, regardless of flight duration or time since the last cigarette (*32*). Animal models of other aspects of addictive behavior have been questioned, by us and other authors, because the timing or sequencing of events does not reflect the typical experiences of human drug users (*33*, *34*). The procedure we describe here addresses those concerns in the realm of cue reactivity and its incubation, and is well suited to disentangle the complex array of behavioral and neural mechanisms underlying the contributions of DSs to relapse (*35*–*37*).

## Materials and Methods

### Experimental Design

The goal of this study was to test for the ability of discriminative stimuli signaling cocaine availability to potentiate cocaine-seeking after withdrawal and then determine if this effect would generalize to non-drug rewards. A detailed description of experimental subjects, apparatus and procedures is included in the following subsection. We first provide an overview of the specific behavioral experiments.

#### Experiment 1: Incubation of discriminative stimulus-controlled cocaine-seeking (Figure 1A)

The goal of experiment 1 was to determine the persistence of non-reinforced discriminated cocaine seeking (*relapse* to DS-controlled cocaine-seeking) and to test for the potentiation of this seeking response during abstinence (*incubation* of DS-controlled cocaine seeking). We trained male and female rats using two 3-h daily sessions (morning and afternoon) to press a central retractable lever only during trials in which lever entry was preceded by the illumination of a light stimulus that signaled cocaine (0.75 mg/kg/infusion) availability (DS+ trials) and to suppress responding during trials when availability of the same lever was preceded by a second light stimulus signaling absence of cocaine reward (DS-trials). There were no additional reward-paired discrete cues. After successful training (20 sessions over 10 days), we tested for discriminated cocaine seeking using a within-subjects design, after varying durations of abstinence extending up to 400 days. For relapse testing, we used the same trial-based procedure and recorded the number of successful trials (defined as making at least one lever press during a trial) and the total number of lever presses (recorded separately for DS+ and DS-trials) made during the 3-h sessions. Finally, after complete cessation of cocaine-seeking behavior on abstinence day 400, we assessed the ability of priming injections of cocaine to reinstate DS-controlled cocaine-seeking, using a within-subjects design and an ascending cocaine dose-response procedure.

#### Experiment 2: Abatement of discriminative stimulus-controlled palatable food-seeking (Figure 2A)

The goal of experiment 2 was to determine whether the persistence and potentiation of responding seen in experiment 1 would generalize to a nondrug reward. We first trained male and female rats using 2-h daily sessions (morning or afternoon) to lever press for palatable food reward (45 mg high-carbohydrate pellets) using a procedure similar to the one used in experiment 1. After successful training (16 sessions over 16 days), we tested all rats for discriminated palatable food-seeking using a within-subjects design similar to that in Experiment 1 but using 2-h sessions, after varying durations of abstinence extending up to 200 days.

### Subjects

We used male (n = 16) and female (n = 16) Sprague-Dawley rats (Charles River), weighing 250-350 g prior to surgery and training. In experiment 1 with cocaine self-administration training, we pair-housed rats of the same sex for one week (n=8 each male and female) prior to surgery and individually housed them after intravenous surgery, during training and abstinence phases. In experiment 2 with food self-administration training, we pair-housed rats of the same sex for one week (n=8 each male and female) prior to the start of behavioral training and individually housed them during training and abstinence. For both experiments, we maintained the rats in the animal facility under a reverse 12:12 h light/dark cycle with free access to standard laboratory chow and water in their home cages throughout the experiment. All procedures followed the guidelines outlined in the Guide for the Care and Use of Laboratory Animals (8th edition; http://grants.nih.gov/grants/olaw/Guide-for-the-Care-and-Use-of-Laboratory-Animals.pdf). In experiment 1, fourteen rats successfully completed discrimination training. We excluded one female rat due to catheter patency failure and one male rat due to failure to acquire drug self-administration. Two male rats and one female rat died during the abstinence period. In experiment 2, all sixteen rats successfully completed discrimination training. One male rat died during the abstinence period. For both experiments, we used maximum-likelihood-based multilevel models (SAS Proc Mixed) to account for missing data.

### Drugs

We received 100 mg/ml cocaine-HCl (cocaine) diluted in sterile saline from the NIDA pharmacy. We chose a unit dose of 0.75 mg/kg per infusion for self-administration training based on previous studies (38) and maintained the same unit dose during discrimination training.

### Intravenous Surgery

For experiment 1, we implanted the rats with silastic catheters in their right jugular vein using previously described methods (*17*). We anesthetized the rats with isofluorane gas (5% induction, 1-3% maintenance) and inserted silastic catheters into the jugular vein. We passed the catheters subcutaneously to the mid-scapular region and attached them to modified 22-gauge cannulae cemented in polypropylene mesh (Small Parts) placed under the skin. We administered ketoprofen (2.5 mg/kg, s.c., Butler Schein) after surgery to relieve pain and allowed rats to recover for 5-7 days prior to drug self-administration training. We flushed the catheters daily with sterile saline containing gentamicin (APP Pharmaceuticals, 4.25 mg/ml) during the recovery and training phases.

### Apparatus

We trained and tested all rats in standard Med Associates self-administration chambers (Med Associates ENV-007) enclosed in a ventilated, sound-attenuating cabinet with blacked out windows. Each chamber was equipped with a stainless steel grid floor and two side-walls, each with three modular operant panels. For experiment 1, we equipped the right-side wall of the chamber with a single retractable lever in the center panel, 7.5 cm above the grid floor. We positioned a discriminative stimulus (light, Med Associates ENV-221M) that signaled cocaine availability on the left panel and another discriminative stimulus (light, Med Associates ENV-221M) that signaled unavailability of cocaine on the right panel of the same side wall, equidistant from the central retractable lever and 14.0 cm above the grid floor. In addition to location, we used red or white lens caps to differentiate between the two discriminative cues and counterbalanced them across the 14 boxes used for experiment 1. We connected the rat’s catheter to a liquid swivel (Instech) via polyethylene-50 tubing that was protected by a metal spring and used a 20 mL syringe driven by a single speed syringe pump (Med Associates PHM-100, 3.33 RPM) placed outside the sound-attenuating cabinet to deliver intravenous cocaine infusions. In experiment 2, we used 8 different self-administration chambers. We equipped the left side wall of these chamber with a single retractable lever in the center panel, 7.5 cm above the grid floor. We positioned a discriminative stimulus (light, Med Associates ENV-221M) that signaled availability of palatable food reward on the right panel and another discriminative stimulus (light, Med Associates ENV-221M) that signaled unavailability of food reward on the left panel of the same side-wall, equidistant from the central retractable lever and 14.0 cm above the grid floor. We again used red or white lens caps to differentiate between the two discriminative cues and counterbalanced them across the boxes used for this experiment. We equipped the central panel of the opposite (right) wall with a pellet receptacle (Med Associates ENV-200R2M-6.0) connected to a 45 mg pellet dispenser (Med Associates ENV-203-45) to deliver palatable food-reward.

### Experimental procedures

Experimental timelines for each experiment are shown in Figures 1A and 2A. The self-administration, trial and discrimination training phases for cocaine and food experiments are described separately below. The subsequent abstinence and relapse test phases are the same for both experiments and described together.

#### Cocaine Self-Administration

We trained male and female rats to lever press for cocaine reward during two 3-h sessions per day that were separated by 30-60 minutes. We gave rats Froot Loops (Kellogg Company, USA) in their home cage one day prior to the start of training and then used crushed Froot Loops when necessary to encourage rats to engage with the lever during initial continuous access training. The start of a session was signaled by the illumination of a light cue on the right side of the retractable lever followed 30 s later by the presentation of the central retractable lever for 180 min. The light remained on for the duration of the session and served as a discriminative stimulus for cocaine reward availability. The same light was later used as a discriminative stimulus to signal availability of cocaine during trial-based discrimination training. Throughout the session, responses on this lever were rewarded under a fixed-ratio-1 (FR1) reinforcement schedule and cocaine at a unit dose of 0.75 mg/kg/infusion (0.1 ml/infusion) was delivered over 3.5 seconds. This infusion duration also served as the timeout period, during which lever presses were recorded but not reinforced. It is important to note that the delivery of cocaine was not paired with any discrete cues. At the end of each 3-h session, the discriminative stimulus was turned off and the lever was retracted. We recorded (1) the total number of lever presses and (2) the total number of infusions received during the entire session. We gave rats up to 6 training sessions to acquire stable self-administration responding in the continuous access procedure before switching them to trial training for cocaine reward.

#### Trial training for cocaine reward

We trained rats in two 3-h trial training sessions per day for two days. We gave rats trial training sessions before trial-based discrimination training to (1) habituate rats to the trial format and (b) introduce the two possible trial contingencies separately before we mixed them together during discrimination sessions. The timeline for a single DS trial, and the differences between the two trial types are depicted in Figure 1B. Each session in this phase consisted of 30 discrete trials separated by a variable inter-trial interval – the start of each trial was signaled by the illumination of a discriminative stimulus for 30 s, following which rats were given access to the central retractable lever for 60 s. During this initial trial training, each session consisted of only one of two possible trial types – trials in which cocaine reward was available (DS+ trials) or trials where cocaine reward was not available (DS-trials).

DS+ trials were signaled by the same DS used during continuous access self-administration (light on right side of lever, counterbalanced for red or white light). During DS+ trials, responses on the lever were rewarded under a fixed-ratio-1 (FR1) reinforcement schedule and cocaine reward at a unit dose of 0.75 mg/kg/infusion (0.1 ml/infusion) was delivered over 3.5 s. This infusion duration also served as the timeout period, during which lever presses were recorded but not reinforced. Additional lever presses during this 60 s period were also reinforced on the same schedule. Similar to self-administration training, delivery of cocaine in these trials was not paired with discrete cues. Sixty seconds after lever presentation, the DS+ was turned off and the lever retracted, signaling the end of the trial.

DS-trials were signaled by the other available DS (light on left side of lever, counterbalanced for red or white light). During DS-trials, all responses on the lever were recorded but not reinforced. Sixty seconds after lever presentation, the DS-was turned off and the lever retracted, signaling the end of the trial.

All rats were trained on DS+ trials in the morning session and DS-trials in the afternoon session. We used two behavioral measures to monitor training during this phase – (1) the total number of DS+ vs. DS-trials with at least one lever press and (2) the total number of responses made during DS+ vs. DS-trials during each 3-hour session.

#### Discrimination training for cocaine reward

We trained rats on the trial-based discrimination procedure for two 3-h sessions per day, separated by at least 30 minutes. In each of these sessions, rats received a total of 60 discrete trials; 30 DS+ trials and DS-trials were intermixed and presented in a pseudorandomized order such that rats received no more than two consecutive presentations of the same trial type during the session. Similar to the previous phase of training, we recorded (1) the total number of DS+ vs. DS-trials with at least one lever press and (2) the total number of responses made during DS+ vs. DS-trials during each 3-hour session.

#### Food Self-Administration

We trained male and female rats to lever press for palatable food reward (TestDiet, Catalogue # 1811155, 12.7% fat, 66.7% carbohydrate, and 20.6% protein) during one 1-hour session per day. We gave rats the 45-mg food pellets in their home cage one day prior to the start of training and then used crushed food pellets when necessary to get rats to engage with the lever during initial continuous access training. The start of a session was signaled by the illumination of a light cue on the right of a central retractable lever followed 30 s later by the presentation of the retractable lever for 60 min. The light remained on for the duration of the session and served as a discriminative stimulus for palatable food reward availability. The same light was later used as a discriminative stimulus to signal availability of palatable food reward during trial-based discrimination training. Throughout the session, responses on this lever were rewarded under a fixed-ratio-1 (FR1) reinforcement schedule. Successful completion of the FR requirement led to the delivery of three 45-mg ‘preferred’ or palatable food pellets over 3.5 s. This reward delivery duration was not paired with any discrete cues and served as the timeout period, during which lever presses were recorded but not reinforced. At the end of the 1-h session, the discriminative stimulus was turned off and the lever was retracted. We recorded (1) the total number of lever presses and (2) the total number of rewards received during the entire session. We gave rats up to 3 training sessions to acquire stable self-administration responding before switching them to trial training for palatable food reward.

#### Trial training for palatable food reward

We trained rats in two 1-hour trial training sessions in two days. We gave rats two trial training sessions before trial-based discrimination training to (1) habituate rats to the trial format and (2) introduce the two possible trial contingencies separately before we mixed them together during discrimination sessions. The timeline for a single DS trial, and the differences between the two trial types are depicted in Figure 2B. Each session in this phase consisted of 30 discrete trials separated by a variable inter-trial interval – the start of each trial was signaled by the illumination of a discriminative stimulus for 30 s, following which rats were given access to the central retractable lever for 60 s. During this initial trial training, each session consisted of only one of two possible trial types – trials in which palatable food-reward was available (DS+ trials) or trials where no palatable food reward was available (DS-trials). DS+ trials were signaled by the same DS used during continuous access self-administration (light on right side of lever, counterbalanced for red or white light). During DS+ trials, responses on the lever were rewarded under a fixed-ratio-1 (FR1) reinforcement schedule. FR completion resulted the delivery of a single 45-mg palatable food pellet after 1 s and a 3.5 s timeout period during which lever presses were recorded but not reinforced. Additional lever presses during this 60-s period were also reinforced on the same schedule. Similar to self-administration training, delivery of food-reward in these trials was not paired with discrete cues. Sixty seconds after lever presentation, the DS+ was turned off and the lever retracted, signaling the end of the trial.

DS-trials were signaled by the other available DS (light on left side of lever, counterbalanced for red or white light). During DS-trials, all responses on the lever were recorded but not reinforced. Sixty seconds after lever presentation, the DS-was turned off and the lever retracted, signaling the end of the trial.

All rats were trained on DS+ trials in the morning session and DS-trials in the afternoon session. We used two behavioral measures to monitor training during this phase – (1) the total number of DS+ vs. DS-trials with at least one lever press and (2) the total number of responses made during DS+ vs. DS-trials during each 3-h session.

#### Discrimination training for palatable food reward

We then trained rats on the trial-based discrimination procedure for one 2-h session per day. In each of these sessions, rats received a total of 60 discrete trials; 30 DS+ trials and DS-trials were intermixed and presented in a pseudorandomized order such that rats received no more than two consecutive presentations of the same trial type during the session. Similar to the previous phase of training, we recorded (1) the total number of DS+ vs. DS-trials with at least one lever press and (2) the total number of responses made during DS+ vs. DS-trials during each 3-h session.

#### Abstinence phase

During the abstinence phase for both experiments, we housed rats in individual cages in the animal facility and handled them 1-2 times per week. In experiment 1, after the rats successfully acquired discrimination, we housed them in the vivarium for up to 400 additional days and tested them repeatedly for relapse after progressively longer durations of abstinence from cocaine. In experiment 2, after rats successfully acquired discrimination, we housed them in the vivarium for up to 200 additional days and tested them repeatedly for relapse after progressively longer durations of abstinence from palatable food reward.

#### Relapse test

In experiment 1 and 2, the experimental conditions during relapse tests were the same as the corresponding trial-based discrimination training session, except that responses on the lever were not reinforced in either DS+ or DS-trials (extinction conditions). In experiment 1, infusion pumps were turned off during all relapse tests and all surviving rats were tested 1, 21, 60, 120, 200, 300, and, 400 days after the last discrimination training session. In experiment 2, pellet dispensers were turned off during all relapse tests and all surviving rats were tested 1, 21, 60, 120, and, 200 days after the last discrimination training session. As with discrimination training, we recorded (1) the total number of DS+ vs. DS-trials with at least one lever press and (2) the total number of responses made during DS+ vs. DS-trials during the entire relapse test session. We operationally define the term relapse as the continuation of non-reinforced discriminated drug seeking after a period of abstinence.

#### Cocaine-primed reinstatement test

In experiment 1, after the final relapse test (day 400), we tested the rats (n=11) for cocaine-priming induced reinstatement during four separate sessions, run on consecutive days (days 401-404). On test days for cocaine-priming induced reinstatement, we gave the rats an intraperitoneal (IP) injection of saline or cocaine 10 min prior to the start of the test session. We chose an ascending dose order for cocaine in order (10, 20 mg/kg) to minimize a carry-over effect of a given priming dose on the subsequent priming dose. We tested the same rats for saline-primed reinstatement before and between cocaine-primed reinstatement tests. The experimental conditions during reinstatement test were the same as the trial-based discrimination training session, except that infusion pumps were turned off for the duration of the test and responses on the lever were not reinforced in either DS+ or DS-trials (extinction conditions). As with discrimination training and relapse tests, we recorded (a) the total number of DS+ vs. DS-trials with at least one lever press and (b) the total number of responses made during DS+ vs. DS-trials during the entire cocaine-primed reinstatement test session. The cocaine priming doses were based on previous studies using the reinstatement model (*39*).

### Statistical analyses

As described earlier, not all rats completed all phases of the experiments. In experiment 1, two of the sixteen rats failed to complete discrimination training and were excluded from the study. Three of the fourteen remaining test rats died during abstinence and did not complete all relapse tests. In experiment 2, all sixteen rats completed discrimination training and were tested repeatedly for relapse during abstinence. One test rat died during abstinence and did not complete all relapse tests. Therefore, we used maximum-likelihood-based multilevel models (SAS Proc Mixed) rather than ordinary-least-squares repeated-measures analyses of variance to account for missing data. Both approaches achieve the same objectives, but maximum-likelihood models obviate imputation of missing data and permit more accurate modeling of nonhomogeneity of variance across unevenly spaced time points.

We conducted all statistical analysis on two behavioral measures – (1) the total number of trials of each DS type with at least one lever press (denoted as *“successful” trials*) and (2) the total number of responses made during each DS trial type over the entire session (denoted as *lever presses*). We followed up on statistically significant main effects or interactions with post-hoc tests as described below. Because some of our models yielded multiple main effects and interactions, we report only those that are critical for data interpretation. In preliminary analyses controlling for sex, we saw sex differences in the acquisition of discrimination, but not in the effects of interest (e.g., the intensity of potentiated seeking or time course of incubation of DS-controlled responding). Therefore, we collapsed our analyses across sex for both experiments. Sex-disaggregated data for all phases of each experiment are provided in the supplementary materials (Figures S1, S2), along with the statistical output for all analyses, including those controlling for sex (Tables S1, S2).

In experiment 1, for the analysis of discrimination (Figure 1C, n=14), we used a 2-way factorial model with within-subject factors of discrimination training session (sessions 5-14) and DS type (DS+, DS-), accompanied by Tukey’s Honest Significant Difference (HSD) test where appropriate for pairwise comparisons between DS+ and DS-for each training session. For the repeated relapse tests (Figure 1D, n=11-14), we used mixed 2-way factorial models with the within-subjects factors duration of forced abstinence (1, 21, 60, 120, 200, 300, and 400 days) and DS type (DS+, DS-), followed by Dunnett’s test for pairwise comparisons between the day 1 relapse test and each of the following days’ relapse tests. We also used Tukey’s HSD for pairwise comparisons between DS+ and DS-for each relapse-test day.

For the tests of priming-induced reinstatement (Figure 1E, n=11), we used 2-way models with the within-subjects factors cocaine priming dose (0, 10, 0, 20 mg/kg), and DS type (DS+, DS-). We used Tukey’s HSD for pairwise comparisons between different cocaine priming doses within each DS trial type. We also used Tukey’s HSD for pairwise comparisons between DS+ and DS-for each reinstatement-test day.

In experiment 2, for the analysis of discrimination (Figure 2C, n=16), we used a 2-way factorial model with within-subject factors of discrimination training session (sessions 3-13) and DS type (DS+, DS-), accompanied by Tukey’s HSD test for pairwise comparisons between DS+ and DS-for each training session. For the repeated relapse tests (Figure 2D, n=15-16), we used 2-way factorial models with the within-subjects factors duration of forced abstinence (1, 21, 60, 120, and 200 days) and DS type (DS+, DS-), followed by Dunnett’s tests for pairwise comparisons between the day 1 relapse test and each of the following days’ relapse testing. We also used Tukey’s HSD for pairwise comparisons between DS+ and DS-on each relapse-test day.

In all models, we used a spatial-power error structure to account for autocorrelation across unevenly spaced intervals; this is similar to the use of a Huynh-Feldt or Greenhouse-Geisser correction in a repeated-measures ANOVA. Alpha (significance) level was set at 0.05, twotailed.

## Acknowledgments

This research was conducted in compliance with the guidelines outlined in the Guide for the Care and Use of Laboratory Animals (8th edition; http://grants.nih.gov/grants/olaw/Guide-for-the-Care-and-Use-of-Laboratory-Animals.pdf) and was approved by the Institutional Animal Care and Use Committee and Institutional Biosafety Committee of the Intramural Research Program of the National Institute on Drug Abuse. Funding: This research was supported by the Intramural Research Program of the National Institute on Drug Abuse (BH).

## Author contributions

R.M., B.J.T, J.M.B, Y.S., and B.H. designed the experiments; L.E.K. and J.K.H performed intravenous catheter surgeries. R.M., B.J.T, L.E.K., S.J.W, J.K.H, V.A.L, and J.M.B collected the behavioral data. R.M., B.J.T, L.E.K., S.J.W, J.K.H, V.A.L, J.M.B, Y.S., D.E. and B.H. analyzed the data. R.M., B.J.T, Y.S., D.E. and B.H. wrote the manuscript with feedback from the other authors.

## Data and materials availability

Data needed to evaluate the conclusions in the paper are present in the paper.

## Supplementary Materials

**Figure S1.**
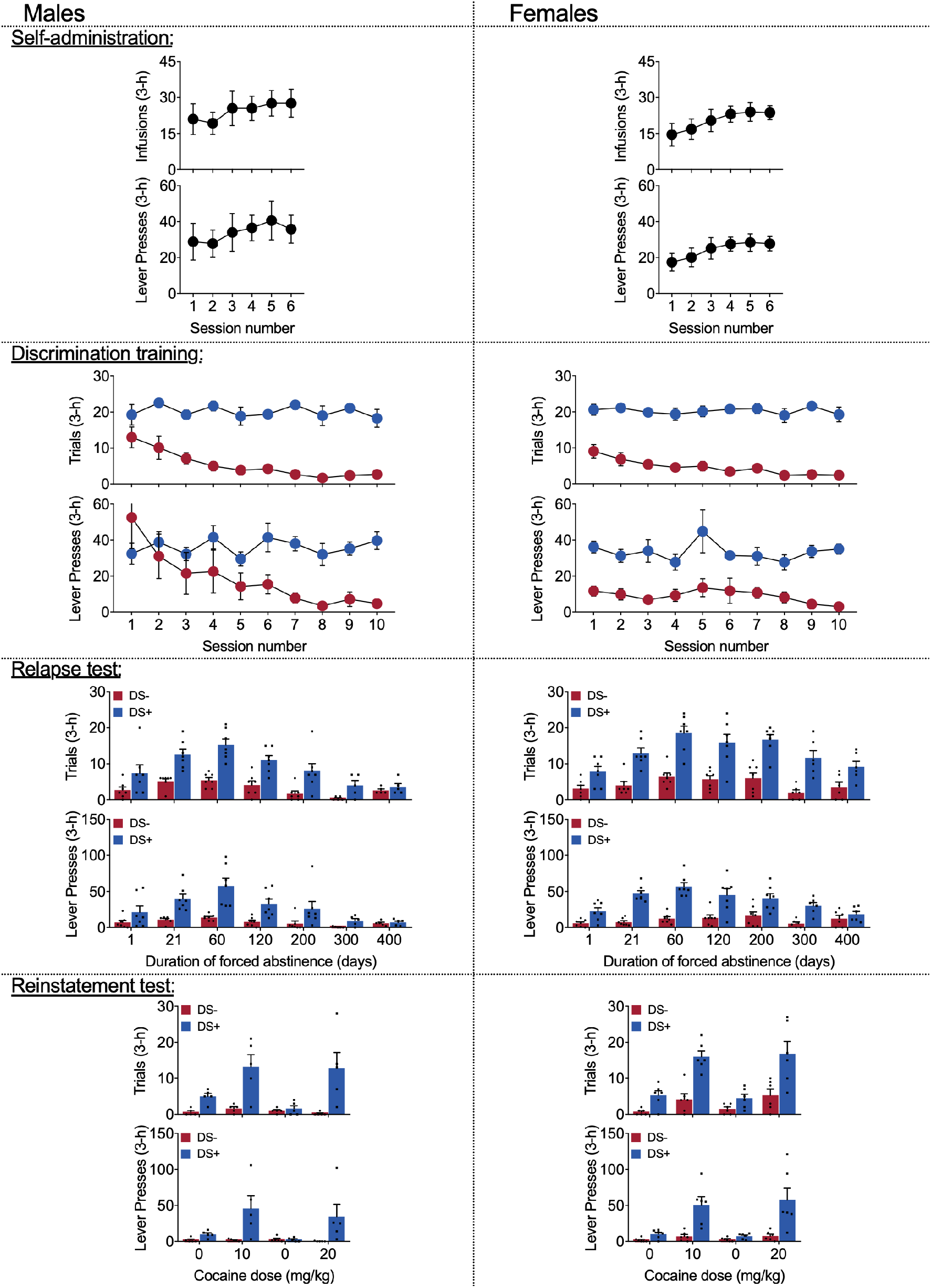
Experiment 1 data disaggregated by sex: Incubation of discriminative-stimulus-controlled cocaine seeking. *<u>Self-administration</u>*: Rats learned to self-administer cocaine over 6 sessions. No main effects or interactions involving Sex were observed. Mean (±SEM) number of cocaine infusions and lever presses during each 3-h session. *<u>Discrimination training:</u>* Over 10 sessions, rats learned to discriminate DS+ from DS-trials. We observed a three-way interaction between DS, Session, and Sex, in both *trials* and *lever presses* measures as male rats pressed more during DS-trials in initial sessions. Mean (±SEM) number of *trials* and *lever presses* during the 3-h discrimination training session. *<u>Relapse test:</u>* Rats showed incubation of lever responding during DS+, but not DS-, trials during abstinence. We observed an interaction between Sex and DS in the *trials* measure but no interaction involving Sex and Day. Further we observed no main effects or interactions involving Sex in the *lever presses* measure. This indicates that while female rats pressed on more DS+ trials than male rats across all test sessions, male and female rats did not differ in the incubation of DS-controlled cocaine-seeking. Mean (±SEM) number of *trials* and *lever presses* during the 3-h relapse test sessions under extinction conditions. Columns indicate mean (±SEM) for the group, while dots indicate values for individual rats. *<u>Reinstatement test:</u>* Rats reinstated DS-controlled cocaine-seeking in response to IP injections of cocaine (10 and 20 mg/kg), but not saline. No main effects or interactions involving Sex were observed. Mean (±SEM) number of *trials* and *lever presses* during the 3-h saline- or cocaine-primed reinstatement test sessions. Columns indicate mean (±SEM) for the group, while dots indicate values for individual rats.

**Figure S2.**
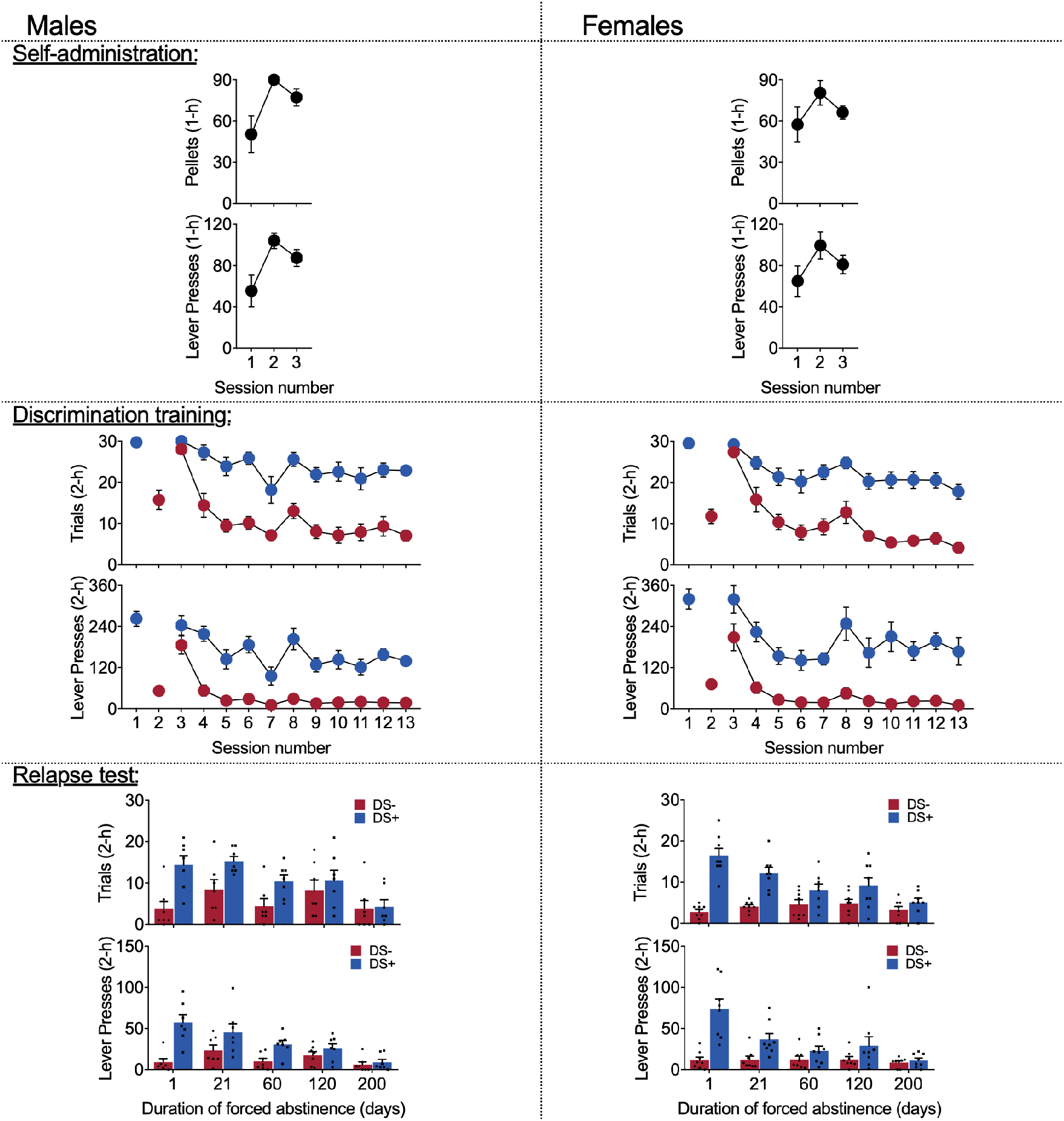
Experiment 2 data disaggregated by sex: Abatement of discriminative stimulus-controlled palatable food-seeking. *<u>Self-administration</u>*: Rats learned to self-administer palatable food pellets over 3 sessions. No main effects or interactions involving Sex were observed. Mean (±SEM) number of cocaine infusions and lever presses during each 1-h session. *<u>Discrimination training:</u>* Over 11 sessions, rats learned to discriminate DS+ from DS-trials. We observed an interaction between Sex and DS in the *lever presses* but not *trials* measure, as female rats pressed more during DS+ trials across all sessions. Mean (±SEM) number of *trials* and *lever presses* during the 2-h discrimination training session. *<u>Relapse test:</u>* Rats’ lever responding during DS+, but not DS-, trials peaked at 1 day of abstinence and abated over 200 days. We observed an interaction between Sex and Day but no interaction involving Sex and DS in the *trials* measure as male rats pressed more during both DS trial types on some days. Further we observed no main effects or interactions involving Sex in the *lever presses* measure. This indicates that while male rats pressed during more trials on some days, male and female rats did not differ in the abatement of DS-controlled palatable food-seeking. Mean (±SEM) number of *trials* and *lever presses* during the 2-h relapse test sessions under extinction conditions. Columns indicate mean (±SEM) for the group, while dots indicate values for individual rats.

**Table S1.**
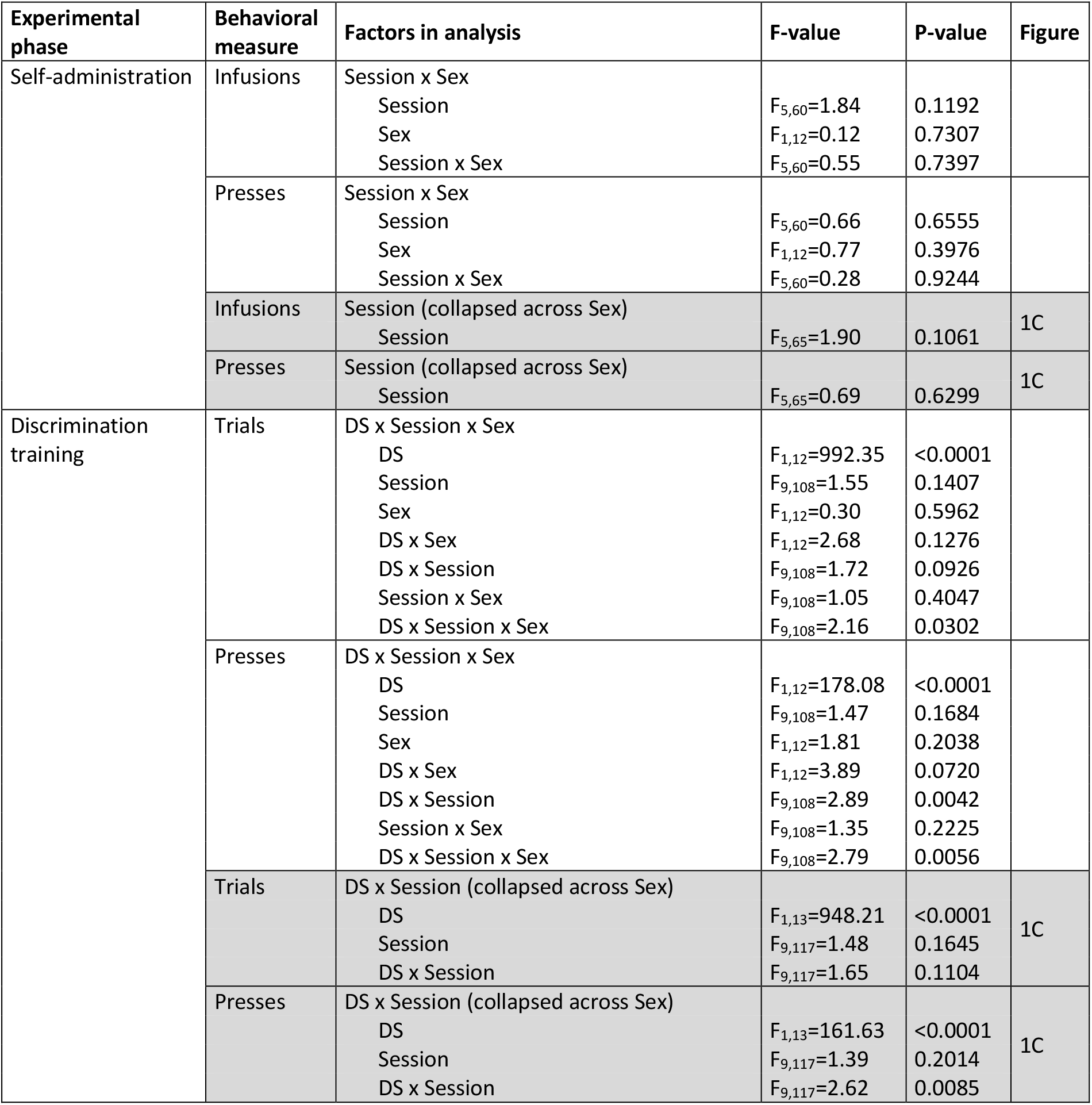
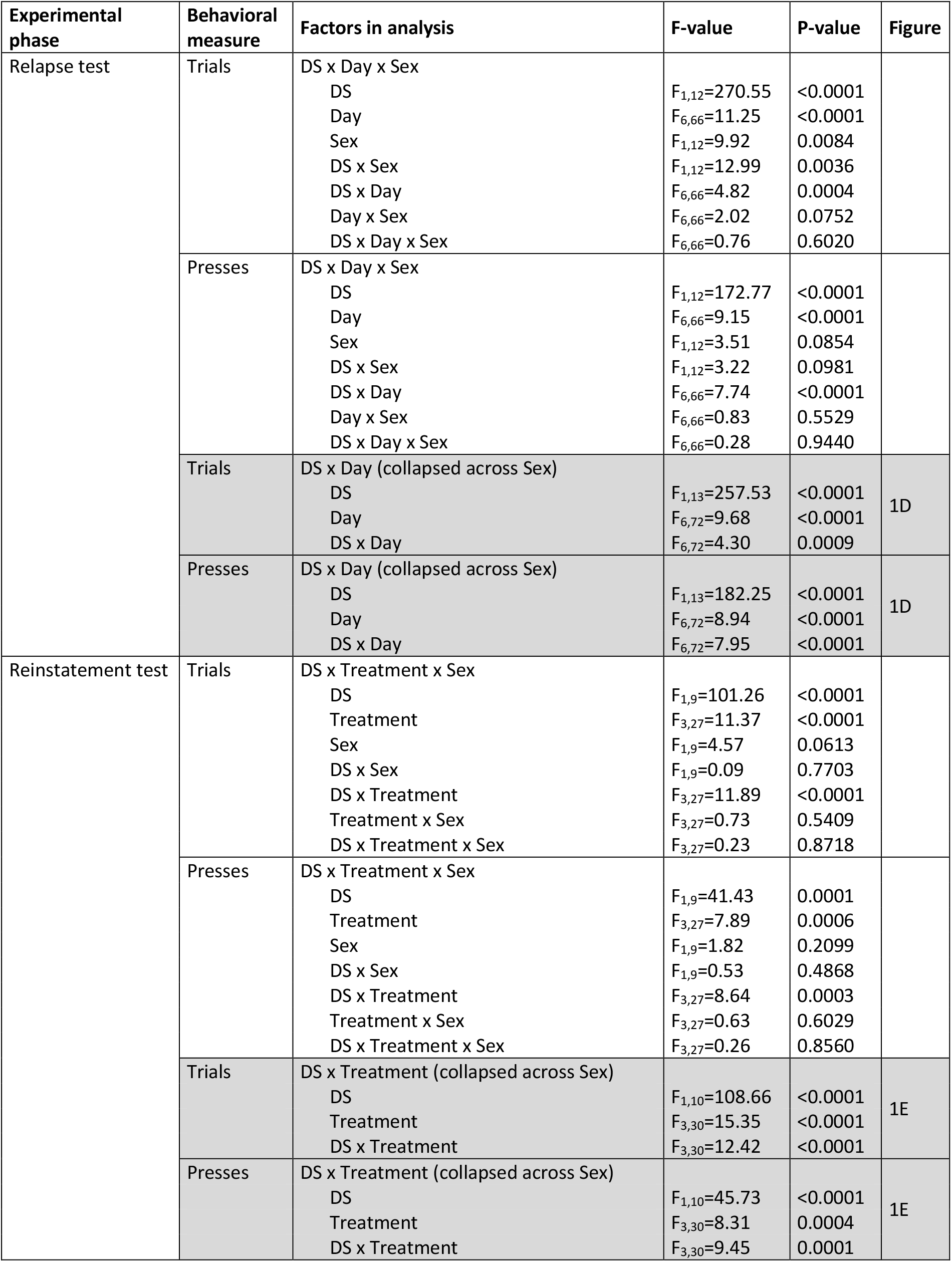
Statistical output for Experiment 1: Incubation of discriminative-stimulus-controlled cocaine seeking (analyses pertaining to figure 1 in the main text are highlighted in grey)

**Table S2.**
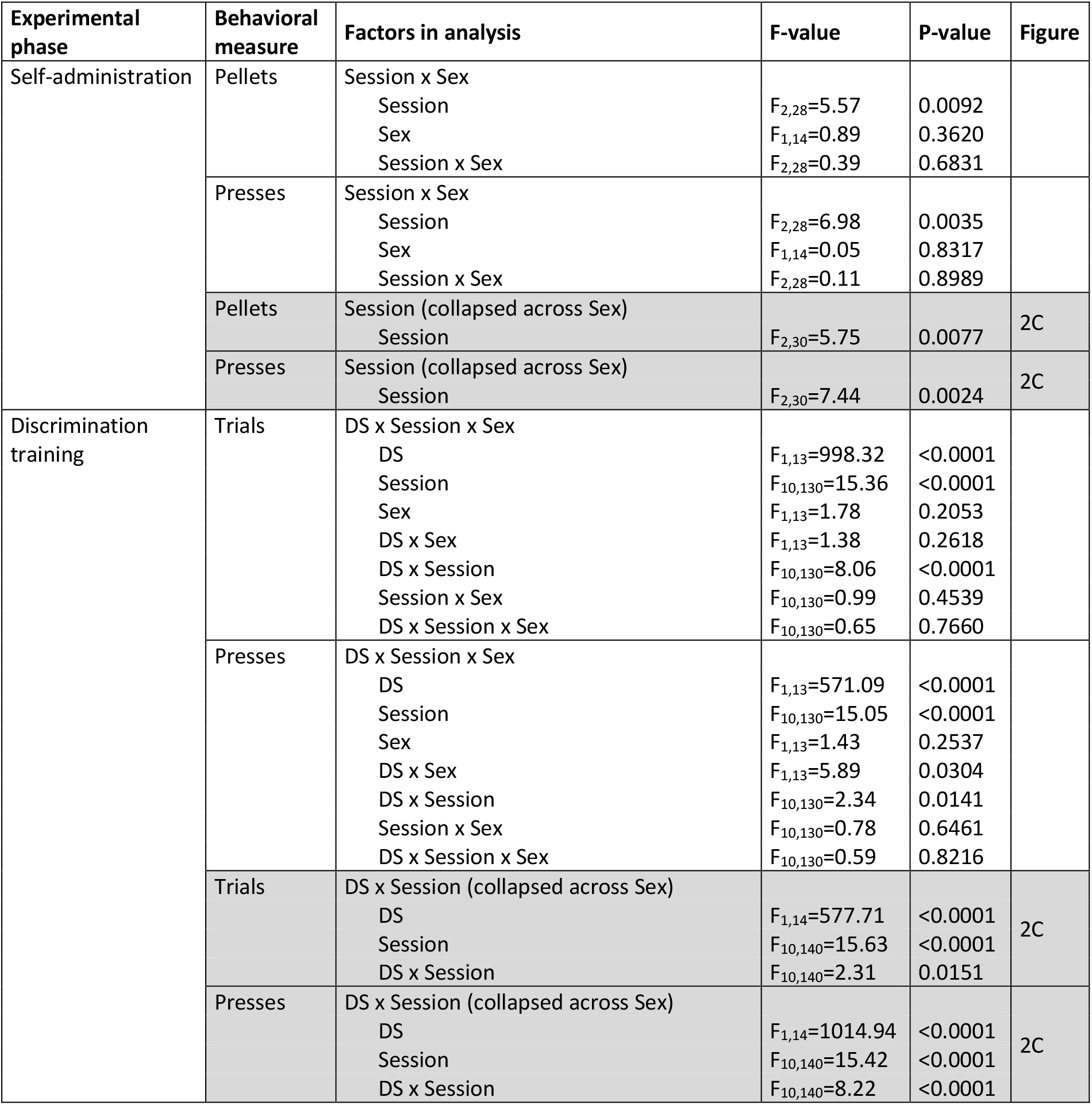
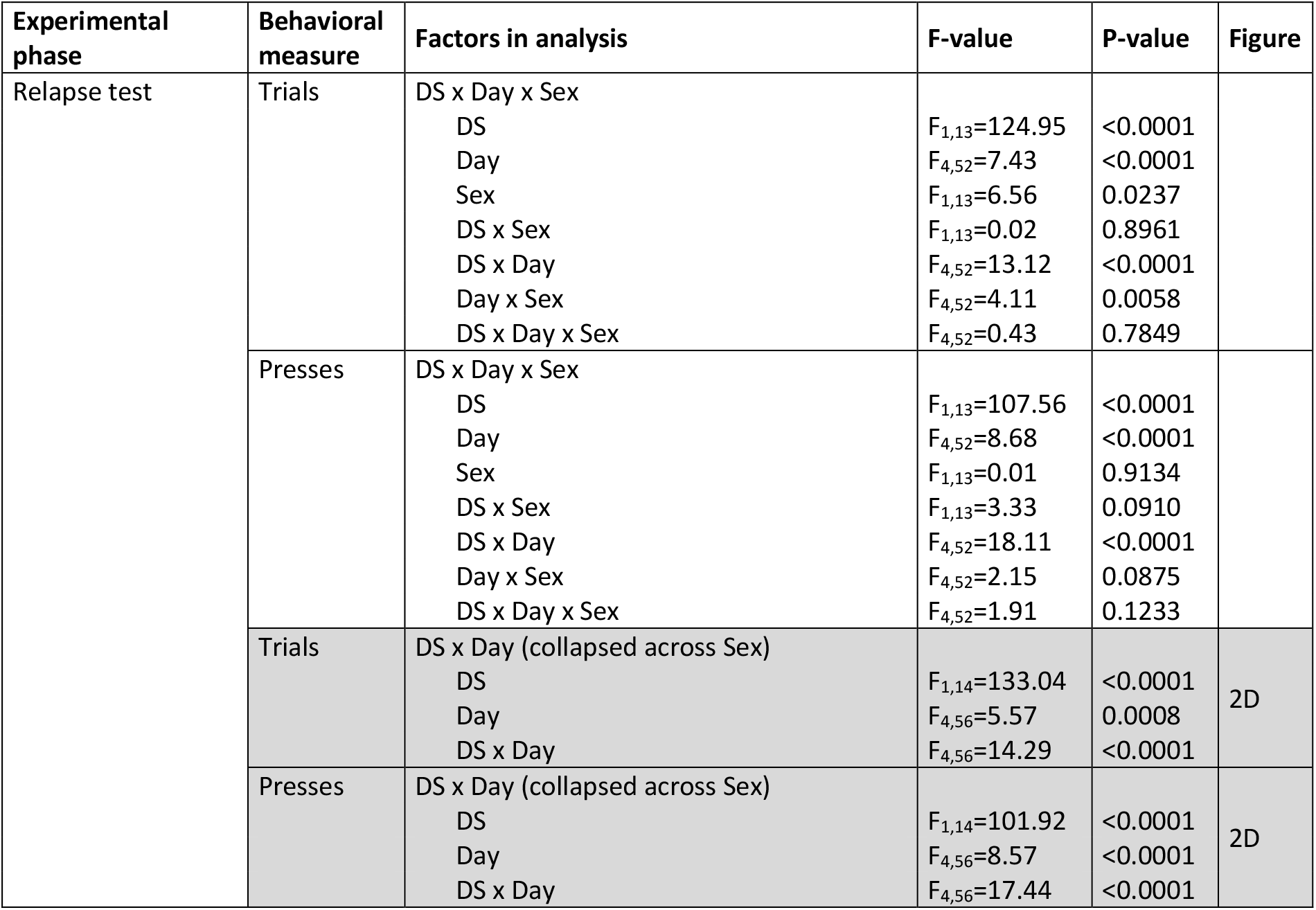
Statistical output for Experiment 2: Abatement of discriminative-stimulus-controlled palatable food-seeking (analyses pertaining to figure 2 in the main text are highlighted in grey)

